# Species interactions promote parallel evolution of global transcriptional regulators in a widespread *Staphylococcus* species

**DOI:** 10.1101/2022.12.19.521106

**Authors:** Casey Cosetta, Brittany Niccum, Nick Kamkari, Michael Dente, Matthew Podniesinski, Benjamin E. Wolfe

## Abstract

Experimental studies of microbial evolution have largely focused on monocultures of model organisms, but most microbes live in communities where interactions with other species may impact rates and modes of evolution. Using the cheese rind model microbial community, we determined how species interactions shape the evolution of the widespread food- and animal-associated bacterium *Staphylococcus xylosus*. We evolved *S. xylosus* for 450 generations alone or in co-culture with one of three microbes: the yeast *Debaryomyces hansenii*, the bacterium *Brevibacterium aurantiacum*, and the mold *Penicillium solitum*. We used the frequency of colony morphology mutants (pigment and colony texture phenotypes) and whole-genome sequencing of isolates to quantify phenotypic and genomic evolution after 15 weeks of the evolution. The yeast *D. hansenii* strongly promoted diversification of *S. xylosus*; by the end of the experiment, all populations co-cultured with the yeast were dominated by pigment and colony morphology mutant phenotypes. Populations of *S. xylosus* grown alone, with *Brevibacterium*, or with *Penicillium* did not evolve novel phenotypic diversity. Whole-genome sequencing of individual mutant isolates across all four treatments revealed numerous unique mutations in the operons for the SigB, Agr, and WalKR global regulators, but only in the *D. hansenii* treatment. Phenotyping and RNA-seq experiments demonstrated that these mutations altered pigment and biofilm production, spreading, stress tolerance, and metabolism of *S. xylosus*. Fitness experiments revealed trade-offs of these mutations across biotic environments caused by antagonistic pleiotropy, where beneficial mutations that evolved in the presence of the yeast *Debaryomyces* had strong negative fitness effects in other biotic environments.

**IMPORTANCE:** Substantial phenotypic and genomic variation exists within microbial species, but the ecological factors that shape this strain diversity are poorly characterized. We demonstrate that the biotic context of a widespread *Staphylococcus* species can impact the evolution of strain diversity. This work demonstrates the potential for microbes in food production environments to rapidly evolve to novel substrates and biotic environments. Our findings may also help understand how other *Staphylococcus* species may evolve in multispecies microbiomes.

## INTRODUCTION

One pattern commonly observed across many microbiomes is morphological, genetic, physiological, and biochemical diversity within microbial species. From metabolic variation in the bacterium *E. coli* (1, 2) to divergent stress responses of the yeast *S. cerevisiae* (3, 4), strain-level diversity is a fundamental feature of most microbes. Intraspecific variation can impact microbial ecology (5–7) and provides useful diversity to exploit for industrial purposes (4, 8, 9). Despite its biological and economic significance, the mechanisms that generate intraspecific phenotypic variation within microbiomes are poorly characterized (10).

Adaptation to the challenges and opportunities presented by other microbial species may contribute to the generation of microbial strain diversity. Due to their large population sizes and fast generation times, microbes can rapidly adapt to a range of selective pressures (11–15). Microbiologists have used experimental evolution to repeatedly subculture a microbial species over many generations and monitor rapid adaptation to novel environments (11, 13, 16). These experiments typically use monocultures of model species in the lab (17–19), but most microbes in nature live in multispecies communities where they experience strong selective pressures from other microbes (20–25). Because biotic interactions have been largely excluded from experimental evolution, our understanding of the dynamics and mechanisms of microbial adaptation in multispecies microbiomes is extremely limited.

Based on examples from macroorganisms and some preliminary studies in synthetic systems with microbes, biotic interactions can mediate microbial adaptation via reduction of population size and subsequent rates of mutations (26), inducing shifts to an alternative niche via competition (27, 28), production of a novel niche via resource provisioning (29, 30), or by inducing or releasing environmental stresses and impacting selection on stress and defense systems (31). This is not an exhaustive set of all potential mechanisms underlying microbial adaptation to biotic environments and multiple mechanisms may be operating in the presence of a single neighbor.

A few highly synthetic studies of laboratory isolates have started to identify different mechanisms of microbial adaptation to varying biotic environments (28, 30, 32–37). These studies begin to add biological complexity to microbial evolution, but they rarely attempt to explain adaptation of microbial populations in naturally occurring communities where ecological environments of ancestral and evolved strains can be clearly defined. These studies also often use selective environments that may not be relevant to the interactions that microbes experience in naturally forming microbial systems. Moreover, this work has been limited to a few target species (mainly *Pseudomonas fluorescens*) and does not usually identify specific mechanisms underlying microbial adaptation to novel biotic environments (33).

The domestication of wild microbes in fermented food environments provides a unique opportunity to fill major gaps in our understanding of how biotic interactions impact microbial adaptation. In some fermented foods -such as sourdough, kimchi, and certain cheeses - wild microbial species colonize the food substrate and become part of the food microbiome (38–43). As continuous batches of food are made in the same location, recirculating microbial populations can adapt to novel abiotic and biotic environments (43, 44). Hints of adaptive evolution come from comparative studies of domesticated and wild microbes (44–46). But the mechanisms that drive the adaptation of fermented food microbes are unknown due to a limited number of studies experimentally recreating the domestication process (43, 44, 47).

In this work, we use one microbial species that is widespread in cheese rinds - *Staphylococcus xylosus* - as a target species for evolution in different biotic environments. This *Staphylococcus* species is often used as a starter culture in the production of cheese and fermented meats (48), but can also be found associated with various mammals (49). This bacterium is an early colonizer of cheese rinds and interacts with a range of fungal and bacterial species during cheese rind succession (50, **Fig. 1A**). We have observed considerable variation in colony morphology and pigmentation of strains isolated from cheese and other fermented food environments (**Fig. 1B**). The underlying drivers of this remarkable phenotypic diversity in *Staphylococcus xylosus* have not been characterized.

**Figure 1.**
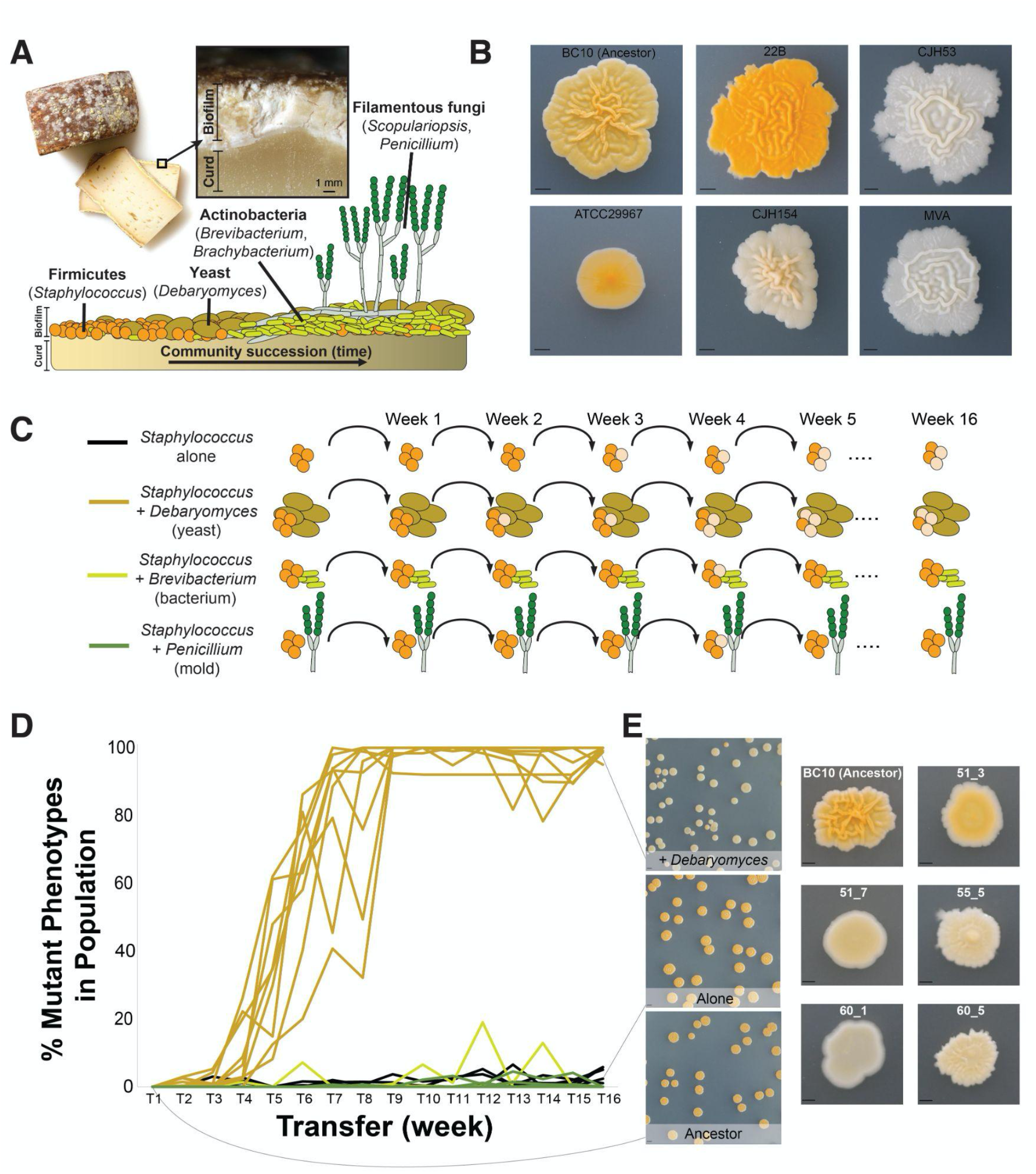
**(A)** Overview of the cheese rind microbiome indicating the main microbial species used in this manuscript including *Staphylococcus, Debaryomyces, Brevibacterium*, and *Penicillium*. **(B)** Phenotypic variation across different isolates of *Staphylococcus xylosus*. All isolates are from cheese except for ATCC29967, the reference isolate for the species. BC10 is the strain used in this study. The scale bar represents 2.5 mm. **(C)** Overview of the experimental design used for the evolution study. **(D)** Frequency of phenotypic mutants in replicate populations across the four experimental treatments. Line colors correspond with the treatments indicated in C. Each line represents a replicate population, with ten replicates per treatment. One +*Debaryomyces* became contaminated and had to be removed from the experiment. Mutant frequency was significantly higher in the *Debaryomyces* treatment compared to the other three treatments at the final transfer (Kruskal-Wallis p < 0.0001, with Mann-Whitney post-hoc tests). **(E)** Examples of typical colony morphotypes within a population of Ancestor (the wild-type BC10), a population evolved Alone (without another microbial species), and a population evolved in the presence of the yeast *Debaryomyces*. Note the distinct difference in color in +*Debaryomyces* compared to the other two treatments. More detailed examples of colony morphologies are shown on the right. 51_3, 51_7, 55_5, 60_1, and 60_5 were used for additional experiments in Figures 3-5. The scale bar in the population photos on the left represents 5mm and the scale bar in the closeup photos represents 1 mm.

To determine if biotic interactions contribute to the phenotypic and genomic diversification of *S. xylosus*, we evolved an isolate of this bacterium (strain BC10) either alone or co-cultured with each of three different cheese rind neighbor species. We predicted that biotic interactions might promote diversification in the presence of some neighboring species that provide novel ecological opportunities and inhibit diversification in the presence of other species that are strong competitors. Because the evolution of colony morphology often correlates with clear underlying adaptive mutations in *Staphylococcus* species (51, 52), we were able to track phenotypic diversity of colonies over time as a proxy for diversification under the different biotic conditions. We also measured genomic diversity at the end of the 16 transfers using whole-genome sequencing of evolved isolates to determine mutations that were driving phenotypic diversity. Competition experiments, phenotypic assays, and RNA-sequencing of select isolates revealed underlying mechanisms of adaptation and intriguing trade-offs associated with the adaptation of *S. xylosus* to different biotic environments.

## RESULTS AND DISCUSSION

### The yeast *Debaryomyces hansenii* promotes phenotypic evolution of *S. xylosus*

We serially transferred populations of *S. xylosus* grown in four different biotic environments: **1)** without a neighbor (hereafter Alone), **2)** with the bacterium *Brevibacterium aurantiacum* (hereafter +*Brevibacterium*), **3)** with the yeast *Debaryomyces hansenii* (hereafter +*Debaryomyces*), and **4)** with the filamentous fungus *Penicillium solitum* (hereafter +*Penicillium*). All of these microbes were isolated from the same natural rind cheese that has been the focus of previous research in this model system, and span a range of life history strategies and temporal dynamics in community succession (50, 53–55). Both *S. xylosus* species and yeasts such as *Debaryomyces* are abundant in early stages of cheese rind development, while *Penicillium* molds and Actinobacteria such as *Brevibacterium* species thrive in the later stages of rind development (50, 54). Because representatives of all three neighboring taxa have been shown to both inhibit or promote the growth of *S. xylosus* and other *Staphylococcus* species (50, 54), we predicted that the different biotic environments created by these neighbors would impact phenotypic and genomic evolution of *S. xylosus*.

Ten replicate populations of *S. xylosus* in the four different conditions were transferred weekly for 16 weeks (about 450 generations). At each transfer, populations were plated out to determine the number of wild-type and mutant colonies. We did not attempt to distinguish different types of mutant colony morphologies within populations because some of these differences are subtle and difficult to track over time. We simply tracked percent of total mutant (not wild-type/ancestor) colonies over time as an approximation of phenotypic evolution within the *S. xylosus* populations. We acknowledge that this approach misses some important dynamics of strain evolution over time and will not capture genomic diversification that does not cause colony morphology changes. But it is a useful tool to track phenotypic diversification that also pointed to some fascinating patterns of genomic evolution at the end of the experiment.

Five weeks into the experiment, we noticed that many colonies across replicate +*Debaryomyces* populations had morphotypes that were noticeably different in appearance compared to the ancestor (**Fig. 1D**). These colonies were often reduced in pigmentation, were not as wrinkly, and often had a shiny appearance that was not apparent in the ancestor (**Fig. 1E**). By the end of the experiment, most of the +*Debaryomyces* replicate populations were taken over by mutant colony phenotypes (99% mean mutant frequency across populations; **Fig. 1D**). In striking contrast to the +*Debaryomyces* treatment, very few mutant colony phenotypes were observed in the Alone (0.02% mutant frequency), +*Brevibacterium* (0% mutant frequency), or +*Penicillium* (0% mutant frequency) treatments.

Differences in population sizes caused by both abiotic and biotic factors can impact rates of evolution (26), and may explain the differences in phenotypic evolution across the four treatments. For example, biotic interactions that suppress population size may inhibit evolution (56, 57). There were significant differences in population size over time across the four treatments, with the +*Penicillium* treatment having the highest mean population size (ANOVA F_3,38_=58.17; p<0.001). However, the +*Debaryomyces* population sizes where we observed the high mutant frequency was not significantly different from the Alone or +*Penicillium* treatments with low mutant frequencies (**Fig. S1**), suggesting that population size does not explain the vastly different amounts of phenotypic evolution we observed.

### Mutations in global regulatory genes explain *S. xylosus* phenotypic evolution with *Debaryomyces hansenii*

To better understand genomic changes that drove the phenotypic diversification of *S. xylosus*, we randomly selected seven colonies from three representative populations from each of the four treatments for whole-genome sequencing. We used Illumina short-read sequencing and mapping to identify SNPs and other genomic changes in each strain. Corroborating the phenotypic evolution observed in the +*Debaryomyces* treatment, we observed a higher frequency of non-synonymous mutations in isolates from that treatment (ANOVA F_3,11_=6.44, p<0.05; **Fig. 2A**). All isolates from the +*Debaryomyces* treatment contained at least one non-synonymous mutation, whereas fewer than half of the isolates from the other treatments had a non-synonymous mutation.

**Figure 2.**
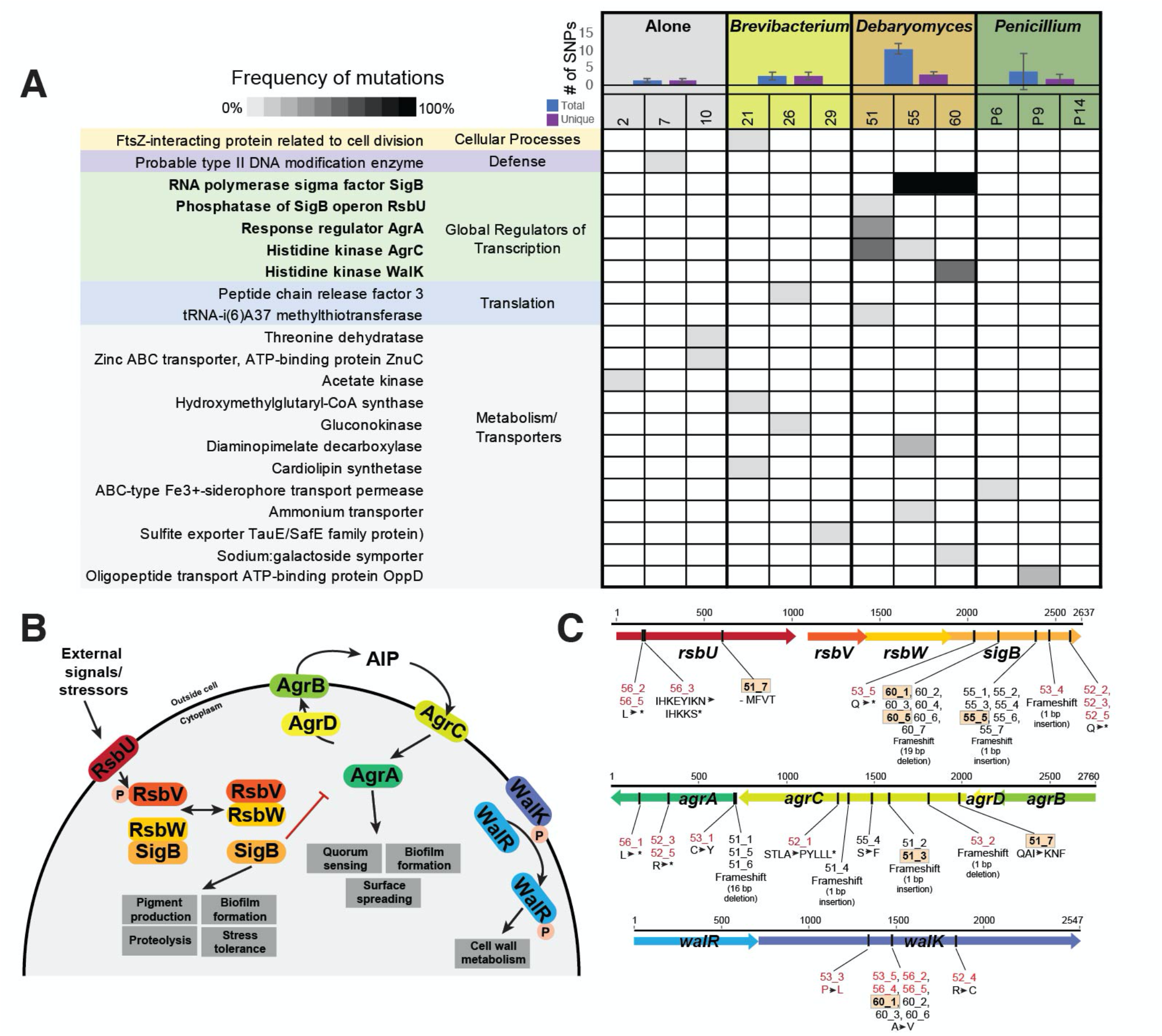
Whole-genome sequencing reveals mutations in putative global regulators of *S. xylosus* BC10 in the +*Debaryomyces* treatment. **(A)** Blue and purple bar graphs show number of SNPs (total in blue, number across unique genes in purple) detected across *S. xylosus* from three populations within each of the four biotic treatments (Alone, +*Brevibacterium*, +*Debaryomyces*, and +*Penicillium*). Shaded cells show frequency of non-synonymous mutations within a gene within a population (columns) based on sequencing 7 isolates per population. Only genes with a putative function assigned are shown. See Fig. S2 for a full table of all mutations, including those with unknown functions and those in the additional +*Debaryomyces* treatments that were sequenced. Numbers 1-10, 21-30, 51-60, and 61-70 are used to indicate unique populations. There were other experimental populations included in the initial experiment (31-40, 41-50), but they are excluded from this manuscript because the neighbor treatment went extinct. **(B)** Overview of the components of the three global regulatory systems where many mutations were detected in the +*Debaryomyces* treatment. These systems are poorly characterized in *S. xylosus* so the potential structure and function of these systems is inferred from what is known in S. aureus. **(C)** Location and type of mutations observed in the sigB, agr, and *walKR* loci. Strains indicated in red font were from the additional +*Debaryomyces* treatments that were sequenced and are not included in A, but are found in Fig. S2.

When considering the types of genes with mutations across all treatments, a very striking pattern emerged: all isolates from the +*Debaryomyces* treatment had at least one mutation in a global regulator of transcription including the alternative sigma factor B (SigB), the accessory gene regulator (Agr), and the WalKR system (**Fig. 2A**). As discussed below, all three of these systems regulate key cellular processes, including biofilm and pigment formation, and could explain the expansion of novel colony morphotypes in the +*Debaryomyces* treatment. We did not detect mutations in global regulators of transcription in any of the other treatments. Mutations in genes related to metabolism and nutrient transport, translation, defense, and other cellular processes occurred across all four treatments, but there was not a specific enrichment of these mutations in any treatment (**Fig. 2A**). To confirm that this pattern of mutations in global regulators was consistent beyond the three +*Debaryomyces* populations that we sampled (51, 55, and 60), we sequenced genomes of 5 isolates from an additional three populations (52, 53, 56) from the +*Debaryomyces* treatment and found an identical pattern of mutations in global regulators of transcription across all three populations (**Fig. S2**).

SigB is a very well-characterized operon in *S. aureus* and other Gram-positive bacterial species where it has been shown to regulate the expression of hundreds of genes (58–60)(**Fig. 2B**). The *sigB* operon includes the genes *sigB, rsbU, rsbV*, and *rsbW* (**Fig. 2C**). A diverse set of mutations in the sigB operon were detected across populations, with most mutations observed in *sigB*, a few in *rsbU*, and no mutations were observed in *rsbV* or *rsbW* (**Fig. 2C**). Mutations in *sigB* ranged from amino acid substitutions to both small (1 bp insertion) and large (19 bp deletion) frameshift mutations. Some populations contained the same mutation across all isolates (e.g. a major deletion of 19 base pairs in all isolates from population 60; **Fig. 2C**) whereas other populations had multiple strains with unique *sigB* mutations (e.g. a predicted truncation in 53_5 and a 1bp insertion causing a frameshift 53_4). Three different mutations were detected in *rsbU*:

Agr is a quorum sensing regulator that has been well-characterized in *S. aureus* and other *Staphylococcus* species (61–63)(**Fig. 2B**). The *agr* operon consists of *agrA* and *agr*C (encoding a two-component signal transduction system), *agrD* (encoding a propeptide that becomes the quorum sensing molecule), and *agrB* (encoding a transmembrane protein). Consistent with patterns of Agr evolution in *S. aureus* (62), all mutations we observed were in *agrA* and *agrC* (**Fig. 2C**). Most mutations were observed in population 51, with four unique mutations distributed across the 7 isolates, including both large deletions and small insertions causing frameshifts (**Fig. 2C**). An amino acid substitution was observed in one isolate from population 55. Additional *agrA* and *agrC* mutations were observed in the additional +*Debaryomyces* populations that were sequenced (**Fig. 2C; Fig. S2**).

The *walRK* operon (sometimes referred to as *yycGF* or *vicKR*) is another regulatory region where multiple mutations were observed in +*Debaryomyces* isolates. The WalKR system is a two-component system that controls a variety of processes in *S. aureus*, including cell wall metabolism (64)(**Fig. 2B**). Although the functions of WalKR have not been characterized in *S. xylosus*, previous work in *S. aureus* has demonstrated that single mutations in *walK* or *walR* can have dramatic impacts on drug resistance and cell wall structure (65). Three different amino acid substitutions were observed in the predicted *walK* gene, including one mutation that occurred across three different populations (53, 56, and 60; **Fig. 2C**). Some strains had mutations in both the *walK* and either *rsbU* (strains 56_2, 56_5) or *sigB* (strains 53_5, 60_1, 60_2, 60_3, 60_6).

Collectively, our resequencing of evolved mutants highlights parallel mutations in global regulators across replicate +*Debaryomyces* populations. The regulatory systems noted above have not been characterized in *S. xylosus*, so we cannot know for sure that they regulate the same genes and traits as in *S. aureus* where they have been well-studied. But many components of these systems are conserved across *Staphylococcus* species or Gram-positive bacteria more generally (66–68), and may operate in a similar manner in *S. xylosus*.

### *S. xylosus* mutants have altered pigment, biofilm, and stress tolerance phenotypes

Because whole-genome sequence data identified mutations in global regulators known to control phenotypes in *Staphylococcus* species, we used a series of assays to characterize these traits in a subset of wild-type and mutant strains of *S. xylosus* BC10. Our goal with these assays was to better understand how the mutations impacted the biology of *S. xylosus* and whether the mutations in the global regulators had similar effects in *S. xylosus* as they do in other *Staphylococcus* species. We selected mutants from different replicate populations of the +*Debaryomyces* treatment that spanned a range of genes and different mutations within those genes (strains 51_3, 51_7, 55_5, 60_1, and 60_5). These selected mutants did not have other mutations in predicted coding regions, so changes in phenotypes observed in these strains could be attributed to mutations in the global regulators.

Colonies of *S. xylosus* evolved in the +*Debaryomyces* treatment had noticeably lighter colonies compared to other populations (**Fig. 3A**). We predicted that these differences could be due to changes in production of the carotenoid staphyloxanthin. This pigment gives *S. aureus* and other *Staphylococcus* species a golden or orange appearance and is encoded by the *crtOPQMN* operon that is controlled by SigB (69). Mutations in the SigB operon can lead to reduced pigmentation (52, 70). Colorimetric assays indicated that the ancestor produced significantly more staphyloxanthin pigment compared to all other mutants (one-way ANOVA; p<0.001). The strains containing a mutation in the *sigB* system produced the least amount of pigment, with strain 60_5 only producing 43% compared to the ancestor. Strains 51_3 and 51_7 also produced significantly less pigment, 68% and 60% compared to the ancestor, respectively.

**Figure 3.**
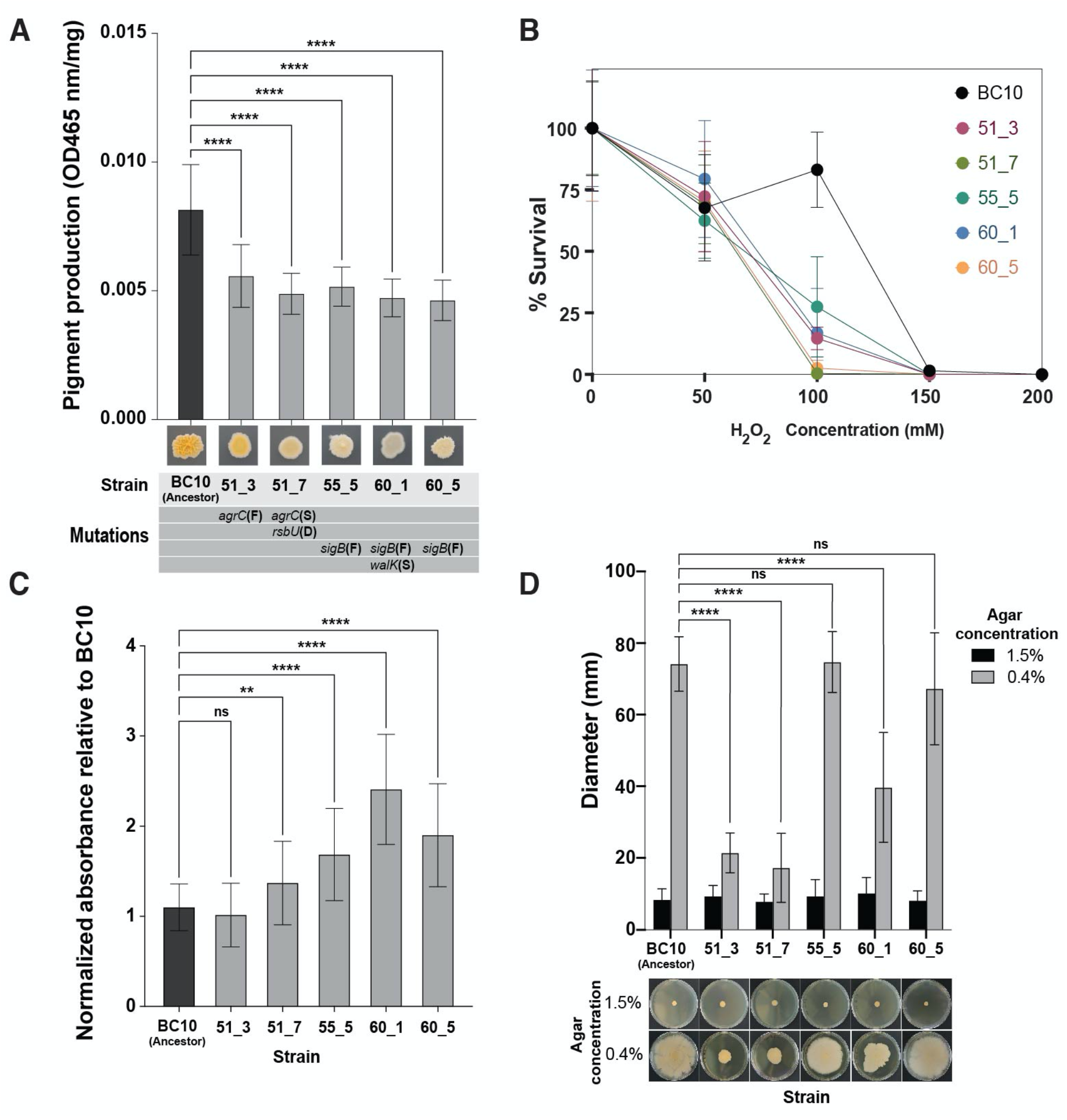
Phenotypic assays reveal biological impacts of global regulator mutations in evolved *S. xylosus* BC10. **(A)** Pigment production in the ancestor and evolved strains of *S. xylosus* BC10. Bars represent means and error bars represent standard deviations. **** indicates *p* < 0.0001 based on ANOVA with Dunnett’s test. N= 3, n= 6. **(B)** Survival of the ancestor and evolved strains of *S. xylosus* BC10 across a range of H_2_O_2_ concentrations. Dots represent means and error bars represent standard deviations. At 100 mM, *S. xylosus* BC10 had a higher survival compared to all mutants (two-way ANOVA, *P* < 0.0001). N= 2, n= 8. **(C)** Biofilm production of ancestor and evolved strains of *S. xylosus* BC10. Bars represent means and error bars represent standard deviations. ** indicates *p* < 0.01 and **** indicates *p* < 0.0001 based on ANOVA with Dunnett’s test. ns = not significant. N= 3, n= 9. **(D)** Spreading of ancestor and evolved strains of *S. xylosus* BC10. Photo at bottom of graph shows representative plates from the 1.5% (low spreading) and 0.4% (high spreading). Bars represent means and error bars represent standard deviations. **** indicates *p* < 0.0001 based on ANOVA with Dunnett’s test. ns = not significant. N= 3, n= 5.

Because carotenoids like staphyloxanthin can function as antioxidants in *Staphylococcus* species (71, 72) and may play a role in how *S. xylosus* interacts with the cheese rind environment, we next tested tolerance of the *S. xylosus* strains to oxidative stress in the form of hydrogen peroxide. All strains were equally susceptible to 50 mM of H_2_O_2_, but the ancestor BC10 had a higher tolerance to 100 mM than the mutant strains (**Fig. 3B**). At 150 mM, there were no viable cells in any population. These data suggest that *S. xylosus* carotenoid levels can provide similar oxidative protection as in *S. aureus* and other Staphylococcus species.

Biofilm formation is a key ecological trait that determines how microbes interact with both abiotic and biotic elements of their environment (73–75). To understand if the observed mutations in the evolved *S. xylosus* affected biofilm production, we used a crystal violet staining assay to quantify biofilm production. In previous studies of *S. aureus*, expression of *sigB* has been linked with the ability to produce biofilms and mutations in *sigB* have resulted in biofilm-negative phenotypes (76, 77). Mutations in the *agr* system can cause increased biofilm production in *S. aureus* because a functional *agr* inhibits biofilm production (78). Based on this, we predicted that *S. xylosus* strains that had mutations in *sigB* would result in decreased biofilm formation compared to the ancestor while the strains with *agr* mutations would have increased or similar levels of production. In contrast to these predictions, strains 55_5, 60_1, and 60_5 (all with *sigB* frameshift mutations), produced significantly more biofilm at 1.53, 2.18, and 1.78 times more than the BC10 ancestor (**Fig. 3C**). Aligning with our predictions, the two strains that had *agrC* mutations (51_3 and 51_7) had increased or equal biofilm formation compared to the ancestor (1.2 times more for 51_7 produced and equal production for 51_3, **Fig. 3C**).

The ability to spread across surfaces has been studied as an important trait for the virulence of *S. aureus*, and may also be important for how *S. xylosus* spreads across cheese surfaces (79, 80). Previous studies have shown the *agr* system has been involved in the production of the biosurfactant that allows this non-motile genera to spread and that disruption of the *agr* system can cause reduced spreading (51, 52, 81). Both *agr* mutants were found to be deficient in colony spreading; strain 51_3 and 51_7 had a 71.1% and 76.7% reduction in spreading compared to the BC10 ancestor, respectively (**Fig. 3D**). Strain 60_1 had an average of 46.5% reduction in spreading with irregular colony edges (**Fig. 3D**). Strains 55_5 and 60_5 were not statistically different from the ancestor (**Fig. 3D**).

These targeted phenotypic data demonstrate that *S. xylosus* mutations that evolved in the presence of the yeast *Debaryomyces* can have major impacts on the biology of this bacterium. Our data also suggest that some of the functions of these global regulators from *S. aureus* may be conserved in *S. xylosus*; predictions about how mutations in the global regulators might alter phenotypes based on *S. aureus* biology were often correct.

### RNA-seq reveals decreased expression of key metabolic pathways in evolved *S. xylosus* mutants

To better understand other underlying changes in the biology of the evolved mutants that were not captured with the phenotypic assays, we used RNA-seq to quantify global transcriptional differences between the ancestor BC10 and three evolved mutants: 51_3, 55_5, and 60_1. We selected these three mutants for RNA-seq because they spanned the spectrum of SigB, Agr, and WalKR and had no or few other genetic changes in other regions of their genome. We compared global gene expression at three days of growth on cheese curd agar because the bacterium is in late exponential phase and RNA can be easily extracted at this time point.

Each of the three evolved strains had major shifts in global transcription compared to the ancestor BC10 (**Fig. 4A-C; Tables S1-S4**). With a differentially expressed gene (DEG) cutoff of log2 ratio of 1/-1 and FDR corrected p-value of 0.05, many genes across the genome of the bacterium had lower expression levels in the mutants: 23% of predicted genes across the genome in 51_3, 13% in 55_5, and 22% in 60_1. A lower, but still substantial number of genes were also upregulated across the three mutants: 18% of genes in 51_3, 7% in 55_5, and 16% in 60_1. These broad patterns of DEGs support previous studies showing that SigB, Agr, and WalKR can have broad transcriptional control in *Staphylococcus* species and that mutations in these global regulators can dramatically alter transcriptional networks within these species (52, 82–86).

**Figure 4.**
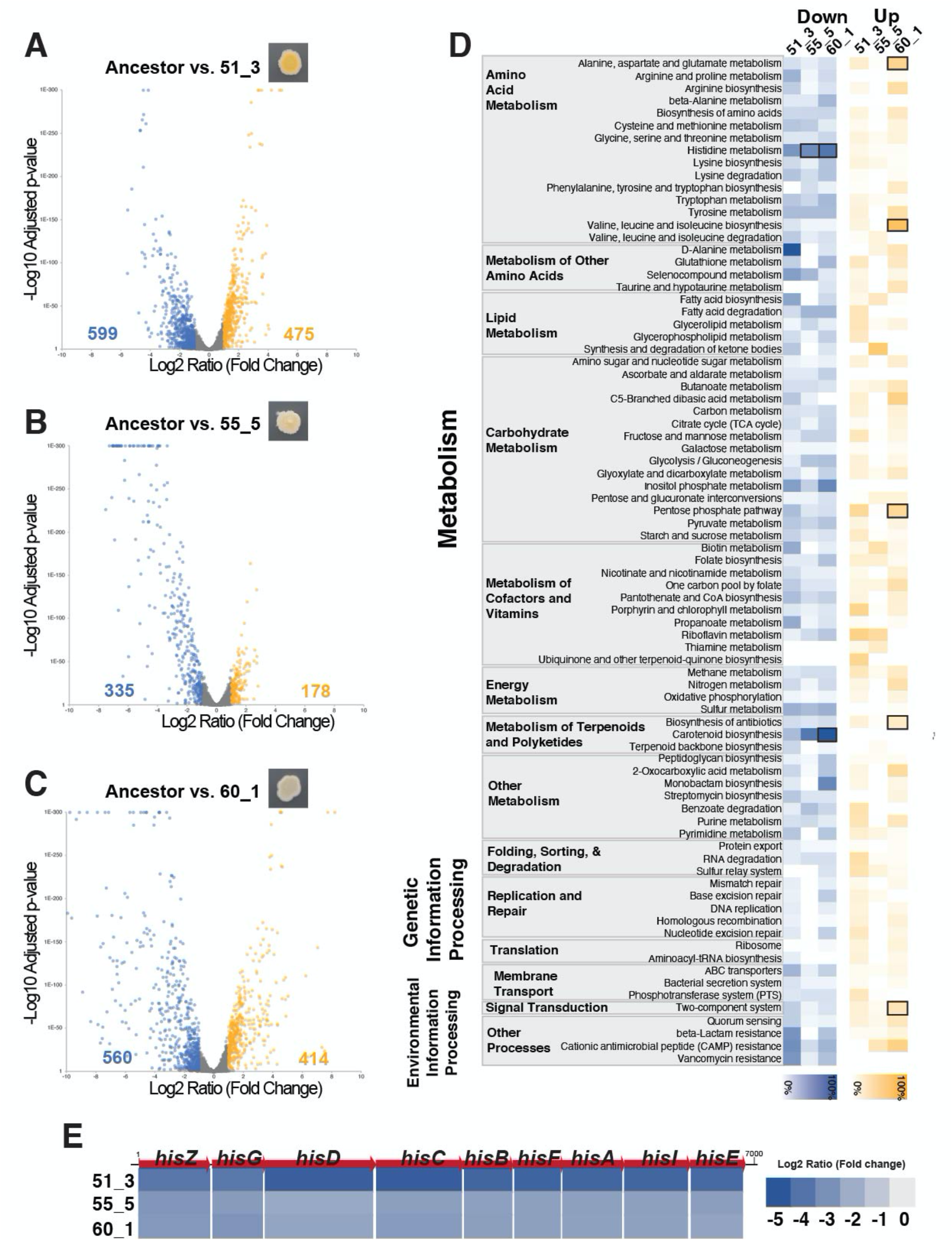
RNA-sequencing reveals transcriptomic impacts of global regulator mutations in *S. xylosus* BC10. **(A-C)** Volcano plots showing changes in gene expression in mutant strain 51_3 (A), 55_5 (B), and 60_1 (C) compared to the ancestor BC10. Each dot represents a gene in the BC10 genome. Yellow dots are genes with significant increases in expression in the mutants. Blue dots are genes with significant decreases in expression compared to the ancestor. Numbers on the left and right of the x-axis indicate the number of genes that were significantly higher (yellow) or lower (blue) in expression for each mutant. **(D)** KOBAS pathway analysis of differentially expressed genes for each mutant. Blue or yellow shading indicates percent of genes in a pathway that were differentially expressed. Bold boxes indicate significant enrichment of a pathway based on a Fisher’s exact test with FDR correction. **(E)** Fold-change in expression of the nine genes in the *his* operon for histidine biosynthesis.

Across the three mutants, most of the differentially expressed genes were associated with metabolism, including the biosynthesis and degradation of amino acids, lipids, carbohydrates,and secondary metabolites (**Fig. 4D; Tables S5-S8**). When we used pathway enrichment analysis to identify what DEGs were significantly associated with all three mutants, two patterns of decreased pathway expression emerged. First, the production of carotenoids, including most of the genes in the staphyloxanthin biosynthesis operon, were downregulated in all three mutants compared to the ancestor (**Fig. 4D**). This pattern was especially pronounced in mutants 55_5 and 60_1 and corroborates our observations and pigment assays above where 55_5 and 60_1 were much lighter in color compared to the ancestor. This also aligns with previous studies that demonstrated that disruption of normal SigB functioning can alter pigment production in *S. aureus* (52).

The other strong pattern of decreased pathway expression across all three mutants was histidine metabolism (**Fig. 4D**). Specifically, all genes in a putative histidine biosynthesis operon were strongly downregulated in all three mutants compared to the ancestor (**Fig. 4E)**. In a previous study of another *S. xylosus* strain, this histidine operon was upregulated in growth in a dairy matrix due to the low availability of histidine and other free amino acids (87). Downregulation of histidine biosynthesis and other amino acid metabolism in the evolved *S. xylosus* BC10 mutants suggests a parallel shift in nutrient requirements in all three mutants, despite different mutations that caused the downregulation.

### Competition experiments reveal fitness trade-offs of evolved mutants in different biotic environments

To better understand how the mutations and altered phenotypes observed above impact the growth and fitness of evolved *S. xylosus* BC10 strains, we conducted a series of assays to measure growth and fitness of all strains in different biotic environments. First, we measured the growth of the ancestor and evolved mutant strains growing individually on cheese curd agar for 7 days. All strains demonstrated typical growth curves on cheese, reaching stationary phase around three days of growth. There were significant differences in growth over time across all strains (F_5,84_ = 6.907, p <0.001; **Table S9**) and this was driven by two major patterns. First, early in growth at day 1, all mutants except 60_5 had lower growth compared to the ancestor (**Fig. 5A**), suggesting a lag in log phase for most mutant strains. Second, at the final time point of 7 days, two strains (55_5, 60_6) had significantly higher abundance compared to the ancestor (**Fig. 5A**). These data demonstrate that the observed genomic changes in the evolved strains had slight, and inconsistent effects on their growth compared to the ancestor.

**Figure 5.**
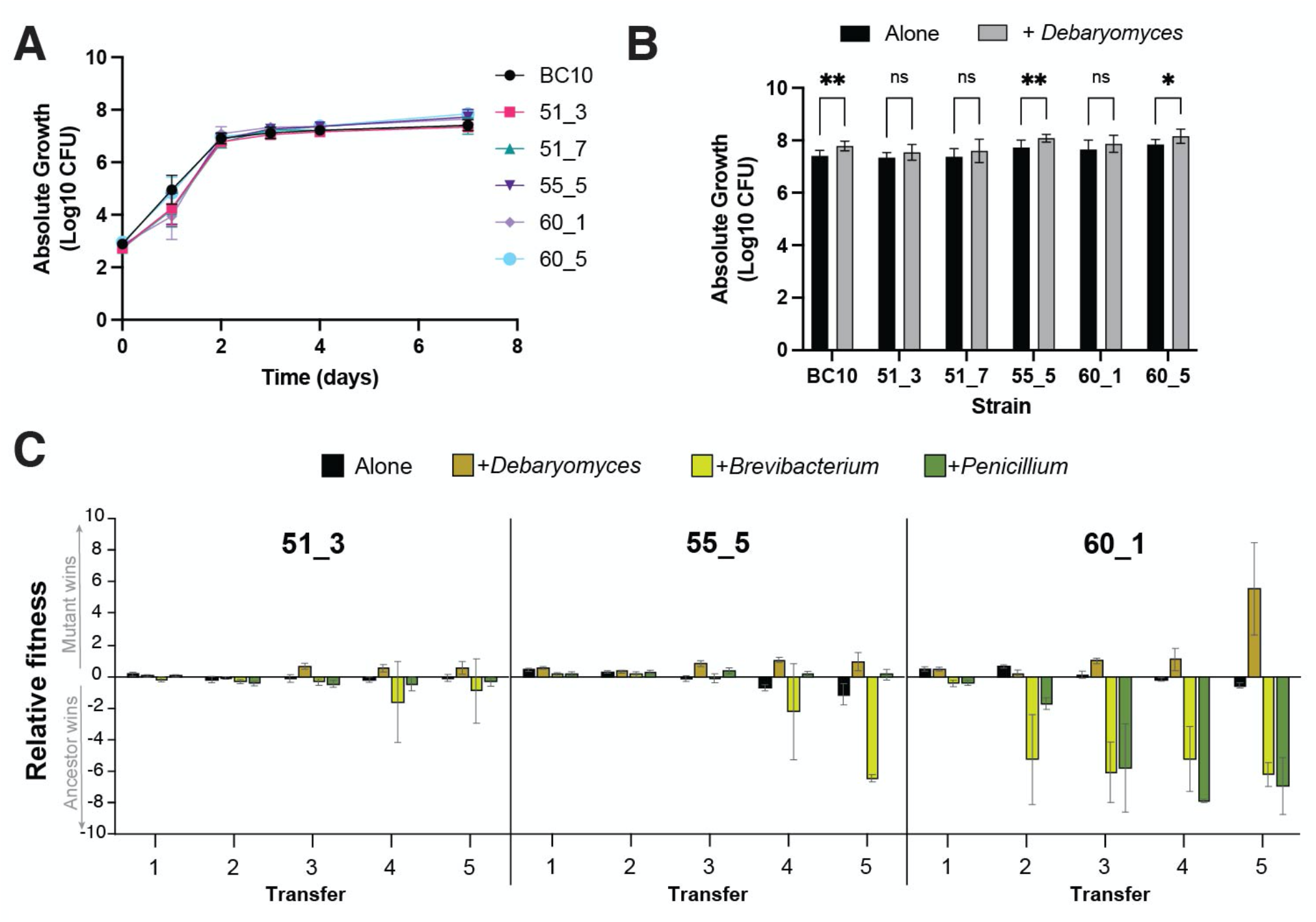
Growth and competition experiments reveal fitness of evolved *S. xylosus* mutants in different biotic environments. **(A)** Growth of ancestor and evolved mutant strains on cheese curd agar alone. Data represent mean CFUs at each time point and error bars are standard errors of the mean. N=3, n=5. **(B)** Growth of ancestor and evolved mutant strains on cheese curd agar with and without the yeast *Debaryomyces* after 7 days of growth. Data represent mean CFUs and error bars are one standard error of the mean. N=3, n=5. *Debaryomyces* increased the growth of the isolates compared to growth alone (F_5,161_= 3.406; P < 0.001). For post-hoc comparisons, ** = p<0.005; * = p<0.05. **(C)** Fitness of ancestor and mutant strains of *S. xylosus* when competed in initially identical ratios in different biotic environments. The ancestor:mutant mix was grown in four treatments and passaged five times to mimic the repeated cycles of growth in the evolution experiment. Relative fitness is expressed as log10((CFUs of mutant strain +1) ÷ (CFUs of ancestor strain +1)). A positive relative fitness means that the mutant strain reached a higher proportion of the total number of CFUs when competing with the ancestor. A negative fitness means the ancestor strain reached a higher proportion. Error bars represent one standard deviation of the mean. n = 8.

Given the high frequency of global regulator mutants in the +*Debaryomyces* treatment, we next tested how the presence of *Debaryomyces* impacted the growth of each of the ancestor and mutant strains. We predicted that *Debaryomyces* may increase the growth of mutant strains relative to the ancestor because they should have a fitness benefit from living with the yeast. Surprisingly, all strains were slightly stimulated by the presence of *Debaryomyces*, with significant increases in growth observed for the Ancestor, 55_5, and 60_5 (**Fig. 5B**).

Growth as single isolates does not fully capture mutant fitness in competitive environments; mutants need to compete with ancestor strains during evolution to become dominant lineages at the end of the experiment. Additionally, very small fitness differences that could accumulate to have major impacts over time may not be apparent in the short time scales used in the previous experiments. Therefore, we next conducted fitness experiments where we grew each mutant in a 50:50 ratio with the ancestor BC10 strain. Because the mutants had unique colony morphologies, we were able to directly compete the strains and distinguish them on output plates. The ancestor:mutant pairing was grown either alone (without any other neighboring microbes) or in the presence of the biotic treatments used in the evolution experiment (+*Debaryomyces*, +*Brevibacterium*, and +*Penicillium*). We transferred the competition experiments five times after seven days of growth to mimic the passaging that occurred in the original evolution experiment. We predicted that mutants would have a higher fitness relative to ancestor strains in the +*Debaryomyces* given how they dominated those populations in the evolution experiment. We only did these experiments with the three mutants used in the RNA-seq analysis (51_3, 55_5, and 60_1) because they spanned the range of phenotypes and mutations observed in the evolution experiment.

As predicted, mutants 51_3, 55_5, and 60_1 all had a significantly higher relative fitness (were a higher % of the total bacterial population compared to the ancestor) in the +*Debaryomyces* treatment after multiple rounds of passaging (ANOVA 51_3 F_3,29_=7.92, *p*<0.001; 55_5 F_3,29_=8.51 *p*<0.001; 60_1 F_3,29_=10.19 *p*<0.001; **Fig. 5C**). In the Alone treatment with no neighbors, all mutants had slight negative fitness compared to the ancestor, which corroborates the lack of these mutants in the Alone evolution populations. We repeated this experiment again with the Alone and +*Debaryomyces* treatment and observed the same pattern of increased fitness of the evolved mutants in the +*Debaryomyces* treatment and lower mutant fitness in the Alone treatment (**Fig. S3**).

Some mutants had striking fitness defects in other biotic environments, especially 60_1 in the +*Brevibacterium* and +*Penicillium* environments (**Fig. 5C**). This suggests that the altered SigB, Agr, and WalKR regulation in these mutants impairs their growth relative to the ancestor strain and explains why they did not evolve in the Alone, +*Brevibacterium*, or +*Penicillium* treatment. We do not know the exact impact of the mutations on WalKR functions in 60_1, but this regulator is known to play key roles in cell wall metabolism (86) and altered functions of WalKR may make mutants unable to grow well in the presence of +*Brevibacterium* or +*Penicillium*. The severe fitness tradeoffs observed for the mutants in different biotic environments is a clear example of antagonistic pleiotropy (88), where beneficial mutations in the presence of the yeast *Debaryomyces* can have strong negative fitness effects in other biotic environments.

## CONCLUSION

One of the most striking overall findings from this work is that interspecific microbe-microbe interactions can shape the evolutionary trajectories of bacteria. While in the Alone, +*Brevibacterium*, and +*Penicillium* treatments we did not see considerable phenotypic evolution or consistent mutations in populations, the yeast *Debaryomyces hansenii* promoted genomic and phenotypic diversification of *S. xylosus*. In a short period of time, many different mutant colony types emerged that were distinct from the ancestral colony morphology, with unique pigmentation and colony structure. This pattern emerged across replicate populations and involved distinct mutations in the same global regulator loci, providing strong evidence of parallel adaptation in this biotic environment.

There are several key nuances and limitations of our work that should be considered when contextualizing it within the cheese rind microbiome and other microbial systems. First, we did not attempt to prevent or control the evolution of the neighbors in our experiment, so the selection pressures experienced by *S. xylosus* throughout time may have shifted if ecologically relevant traits of the neighbors evolved. This was a conscious decision as we were trying to mimic population dynamics in cheese aging facilities where both target microbes and neighboring microbes might evolve. We did not observe any obvious growth or colony morphology changes in the neighbors and previous research in our lab has demonstrated that one neighbor (*Penicillium*) does not evolve new phenotypes if bacteria are present (47). Second, our studies were confined to a controlled laboratory environment, so we do not know how often mutants we have observed here would evolve in cheese production and aging environments. However, we believe there is considerable potential for similar evolutionary processes to unfold in cheese aging facilities. In these environments, *S. xylosus* populations experience varying biotic environments due to patchiness of cheese rind communities across the surfaces of cheese and variation in community composition across wheels of cheese (50, 54, 89). Finally, we only considered how one neighboring species at a time impacts the evolution of *S. xylosus and* did not consider how combinations of species drive phenotypic and genomic evolution of this bacterium. For example, combining *Debaryomyces* with a neighbor that decreased the fitness of evolved mutants (e.g. *Penicillium* or *Brevibacterium*) might inhibit the global regulator evolution observed in the *Debaryomyces* alone treatment.

A major question that remains unanswered is why does *Debaryomyces* promote the diversification of *S. xylosus* on cheese? We did not identify specific mechanisms underlying *Staphylococcus*-*Debaryomyces* interactions, but several lines of evidence from our data and the cheese rind system provide some helpful clues. One ecological explanation is that the degree of niche overlap between these two species may promote diversification of *S. xylosus*. A common theme in the study of adaptation to novel environments is that interspecific interactions can promote diversification when there is strong niche overlap (32). *Staphylococcus* species and yeasts such as *Debaryomyces* share similar resource and temporal niches in cheese rind microbiomes. At the early stages of cheese ripening, the pH of cheese is low, salt concentrations are high, and resources are stored away in the casein of cheese. Previous work in our lab and others has shown that both *S. xylosus* and yeasts such as *Debaryomyces* grow early in cheese succession (50, 54, 90). These microbes are both tolerant of the high salt concentrations and have proteolytic abilities that help them access free amino acids in cheese. In contrast, the other two biotic neighbors (*Brevibacterium* and *Penicillium*) grow later in cheese rind succession and occupy different niches compared to *Staphylococcus*. During our evolution experiment, competition for resources between *S. xylosus* and *Debaryomyces* may have promoted the adaptation of new lineages of *S. xylosus* with different metabolic and ecological traits.

Another potential explanation for the rise of mutants with altered global regulation is that these mutants are only able to persist in the presence of *Debaryomyces* due to facilitative interactions between the yeast and mutants. Both the ancestor and evolved *S. xylosus* mutants had slightly higher growth in the presence of *Debaryomyces*, suggesting that the yeast does provide some benefits for the bacterium. RNA-seq demonstrated strong metabolic downregulation of many pathways in evolved mutants, including biosynthesis of histidine and other amino acids. *Debaryomyces* may have promoted the rapid and reproducible rise of these mutants through unknown mechanisms of supplementing their altered metabolism. For example, *Debaryomyces* may break down some cheese curd components and release nutrients that are needed to support the altered metabolism of the *S. xylosus* with mutations in global regulators. Future work using metabolomics will help pinpoint the chemical mediators of *agr* and *sigB* evolution in the *Debaryomyces* environment.

Our evolution experiment demonstrates how biotic interactions might facilitate the generation of strain diversity within microbiomes. These data also contribute to an emerging body of work demonstrating that microbes have very different evolutionary outcomes when biotic complexity is incorporated into experimental evolution (30, 33, 35, 36, 91, 92). Because most medically and industrially important microbes often live with other microbial species that may impose strong biotic selection on target microbial species (23, 25, 93, 94), management and engineering of microbiomes may need to incorporate how microbe-microbe interactions can shape evolution of microbial species. As additional studies use a broader range of target microbes and more diverse biotic environment treatments, we can begin to create a predictive framework for when and how biotic environments can impact microbial evolution.

## MATERIALS AND METHODS

### Experimental evolution

All yeast and bacterial cultures were maintained as frozen stocks in 15% glycerol in a -80°C freezer. The main target for experimental evolution, *S. xylosus* strain BC10, was originally isolated from the surface of a natural rind cheese made in Vermont, USA and has been previously described (50). Three different microbial strains were used as neighbor treatments: *Brevibacterium aurantiacum* JB5 (as previously described in 53), *Penicillium solitum* strain #12 (as previously described in 47), and the yeast *Debaryomyces hansenii* strain 135B (as previously described in 50). All of these strains were isolated from the same natural rind cheese made in the same aging facility and commonly co-occur in natural rind cheeses made in different regions of the world (50, 54, 95–97).

To initiate the evolution experiments, cultures were inoculated into 1.5 mL microcentrifuge tubes filled with a 150 µL aliquot of cheese curd agar (CCA) containing 3% salt (w/w) (98). Inoculum for each culture came from frozen glycerol stocks where a known CFU/µL concentration of the microbial culture had been previously determined (98). In the alone and +neighbor treatments, 50 CFUs of *S. xylosus* were inoculated into each replicate tube by adding 10 µL of inoculum suspended in 1X phosphate buffered saline (PBS) to the CCA surface. In each of the neighbor treatments, 50 CFUs of a neighbor were added to the *S. xylosus* inoculum. Each treatment was replicated 10 times. One replicate from the +*Debaryomyces* treatment was lost during the experiment due to contamination. Tubes were covered with sterile AeraSeal™ films and then placed in microcentrifuge tube racks containing a snack Ziplock bag with a paper towel moistened with 5 mL of sterile PBS to maintain high humidity. A rack was placed on the tube to contain the moisture and tubes were incubated in the dark for seven days at 24 °C.

After seven days of growth, each population was transferred by adding 300 µL of 15% glycerol to the existing population, homogenizing the sample with a micropestle, and pipetting 10 µL of the homogenate using a wide orifice pipette tip to a fresh 1.5 mL microcentrifuge tube containing 150 µL of cheese curd agar. Each homogenized sample was then frozen at -80 °C to serve as fossil stocks for isolating colonies. This was repeated over a period of 15 weeks.

To determine the frequency of mutant phenotypes within the population at each transfer, 20 µL of homogenate was serially diluted and plated onto plate count agar with milk and salt (PCAMS, (98)) plates with 50 mg/L of cycloheximide to inhibit the growth of fungi. Plates were incubated at 24 °C with natural room light to allow for the development of colony pigmentation. Plates were scored for % mutant frequency by counting colonies and noting the presence of colonies with altered phenotypes. Wild-type *S. xylosus* BC10 is intensely orange/yellow in color, wrinkly, and shiny. Colonies deemed mutants had abnormal pigmentation (white, light yellow), colony texture (less wrinkly to completely smooth), and colony surface appearance (matte or partially shiny). Abundances of neighbors were also determined by plating onto selective media for fungi (PCAMS + 50 mg/L of chloramphenicol to inhibit bacteria) or by counting colonies of *Brevibacterium* that grew alongside the *S. xylosus* colonies.

### Whole-genome sequencing and analysis

To identify genomic changes in evolved isolates, whole-genome sequencing was conducted on seven randomly selected colonies from the final time point (T16) from three populations across the four treatments (84 genomes in total). DNA was extracted from streaks of pure cultures growing on PCAMS plates using a MoBio PowerSoil DNA Isolation kit. Illumina sequencing libraries for each genome were constructed using the NEBNext Ultra DNA Library Prep Kit for Illumina (New England Biolabs) using the manufacturer’s protocol with DNA input concentrations of 5-15 ng/µL, a 20-minute fragmentase incubation for DNA fragmentation, and 5 rounds of PCR enrichment. Equimolar concentrations of each library representing isolate genomes were pooled and sequenced in one run on an Illumina HiSeq 2500 at the Harvard FAS Center for Systems Biology Core Facility. We aimed to have ∼20-30X coverage of each genome for variant detection.

Reads were mapped to a reference genome of *S. xylosus* BC10 (NCBI WGS Project LNPU00000000) using end-to-end alignment in Bowtie2 (99). We also mapped reads of the ancestral isolate to the reference genome to identify any errors in the original genome assembly that could cause erroneous variant calling. Variant calling was conducted using FreeBayes in pooled continuous mode with the following settings: ploidy = 1, minimum alternate count = 10, minimum alternate fraction = 0.9, and minimum probability = 0.1. Any SNPs or other variants identified from mapping raw reads of the ancestor were excluded from variant calls of the evolved isolates. When read mapping indicated potential deletion or other structural variants, we re-assembled the genomes of those isolates using SPAdes and annotated the genome using RASTk.

### Phenotypic assays

To measure staphyloxanthin production, bacteria were streaked across a PCAMS plate and were left to grow for one week to develop pigment. Each plate was laid out singly in a plastic bag to ensure each plate received even exposure to light. After one week of growth at 24°C, 20 mg of cells were removed from the plate with a wooden dowel. Cells were resuspended in 800 µL of methanol and thoroughly mixed via vortexing. The suspensions were left overnight at 55°C with shaking at 525 rpm on a benchtop shaking incubator to extract the pigment from cells. After extracting, the suspensions were centrifuged at 5000 g for 10 minutes. The pellets appeared colorless (white) with the pigment in the supernatant. The absorbance of the supernatant was read at 465 nm using methanol as a blank. The absorbance readings were normalized to cell weight.

To quantify biofilm production, bacteria were grown in 5 mL brain heart infusion (BHI) broth overnight at 24°C. The cells were diluted 100-fold in BHI and 200 µL of inoculum was added to eight wells of a flat-bottom 96-well plate (Falcon). After 16 hours of growth at 24°C, the OD_600_ was recorded to account for differential growth of each strain. Wells were washed three times by manually flipping plates to dispose of liquid and adding 200 µL of 1X PBS to the wells to remove dislodged cells. The cells were heat fixed to the plate by incubating for one hour at 60°C. 50 µL of a solution of 0.06% crystal violet stain was added to each sample. These were incubated for five minutes at room temperature and rinsed three times with 200 µL of 1xPBS. Biofilm formation was estimated by solubilization of crystal violet by adding 200 µL of 30% acetic acid and determining the OD600. For each of three independent experiments, 8-12 replicate wells in three replicate plates were used.

The effect of mutations on bacterial spreading was assessed on soft Brain Heart Infusion (BHI) agar (0.4% w/v). After autoclaving, the medium was cooled down to 50°C and 20 mL was partitioned into Petri dishes. The plates were kept at room temperature for three hours prior to spreading assay. Bacterial strains were grown in 5 mL BHI broth overnight at room temperature. The cells were adjusted to OD_600_ of 1 and 2 µL of the cell suspension was spotted in the center of a soft 0.4% and 1.5% BHI agar plate. The plates were incubated for five days at 24°C. Spreading diameter was calculated by measuring the largest diameter of the colony for each of five replicates.

After 16 hours of growth in BHI broth, inocula of BC10, 51_3, 51_7, 55_5, 60_1, 60_5 were standardized to OD600 of 0.5 and incubated in the indicated concentrations of H_2_O_2_ (0, 50, 100, 150, 200 mM) in the dark at 24°C in a volume of 200 µL. After 45 minutes, the reaction was stopped with the addition of 2 units/mL catalase and incubation for 20 minutes. Cells were serially diluted and plated on PCAMS to quantify CFUs. Values are expressed as a percentage of the CFU in the control lacking H_2_O_2_. Values are the averages of eight replicates in two independent experiments (N=2, n=8).

### RNA-sequencing

To characterize the transcriptomes of the ancestor and three mutants (51_3, 55_5, and 60_1), we constructed and analyzed RNA-seq libraries. Lawns of each strain were grown on 100 mm Petri dishes containing 10% cheese curd agar media with 3% salt. Bacteria were inoculated at 50,000 CFUs and plates were incubated at 24°C for three days. Bacterial cells were harvested from the surface of the cheese agar by using a sterile razor blade and were then immediately placed into 3 mL of RNAProtect Bacteria Reagent (Qiagen) and frozen at -80C until RNA extraction. Total RNA was extracted as previously detailed using a 125:24:1 (v/v/v) phenol/chloroform/isoamyl alcohol extraction (55, 98). Bacterial rRNA was depleted using a NEBNext Bacteria rRNA Depletion Kit and depleted RNA was used to construct libraries using the NEBNext Ultra RNA Library Prep Kit for Illumina. Libraries were sequenced using a NextSeq 550 Illumina Sequencer with single-end 75 base-pair reads. Reads were mapped to the *S. xylosus* BC10 draft genome (GenBank Assembly Accession: GCA_001747745.1) using Bowtie2 and differentially expressed genes were identified using DESeq2 with a log2 ratio of 1/-1 and FDR corrected p-value of <0.05. To identify pathways that were enriched in the differentially expressed genes, we used the Gene-List Enrichment tool of KOBAS (100) with our BC10 draft genome as the background genes to test for enrichment.

### Growth and competition experiments

To determine the ability for the ancestor and evolved mutants to grow on cheese, each was inoculated at a concentration of 667 CFUs onto the surface of 150 µL of cheese curd agar in a 1.5 mL Eppendorf tube. Experimental units were incubated as described for the evolution experiment above. Growth of cells over time was determined at one, two, three, four, and seven days by resuspending the cells into 300 µL of 30% glycerol. The cells and cheese curd were homogenized via pestling, serially diluted, and plated onto PCAMS to determine CFUs. To determine how the yeast *Debaryomyces hansenii* strain 135B impacted the short-term growth of the ancestor and mutant *S. xylosus* strains, the same experimental approach was used with 667 CFUs of *Debaryomyces hansenii* added to the +*Debaryomyces* treatment.

To determine the fitness of mutant strains, we competed three strains (51_3, 55_5, and 60_1) against the ancestor strain either alone (with no neighbor) or in the three biotic treatments (+*Debaryomyces*, +*Brevibacterium*, +*Penicillium*) in 50:50 initial ratios. Each strain of *S. xylosus* (ancestor, mutant, and *Debaryomyces/Brevibacterium/Penicillium* if present) was inoculated at a concentration of 667 CFUs into 1.5 mL Eppendorf tubes containing 150 µL of cheese curd agar. After one week of incubation at 24°C under aerobic conditions, cells were harvested by resuspending the wells into 300 µL of 30% glycerol and homogenized via pestling. Ten µL of the homogenized communities were transferred to a new tube to mimic the experimental evolution and 20 µL was serially diluted and plated onto PCAMS to determine the abundance of ancestor and mutant colonies. Mutant colonies could be distinguished from ancestor colonies due to their reduced pigmentation and altered colony morphology. This experiment continued for a total of five weeks. Relative fitness of the mutant strains was calculated as log_10_((CFUs of mutant strain +1) ÷ (CFUs of ancestor strain +1)). A second competition experiment was conducted to confirm the observed fitness patterns observed in the first experiment. It was executed using the same setup, transfer, and plating methods as above, but with only the alone and +*Debaryomyces* treatments. This second experiment was conducted for only three weeks.

### Statistics

To assess whether there were significant differences in the frequencies of phenotypic mutants or mutations across the experimental evolution populations (Fig. 1 and 2), we used ANOVA with Tukey’s post-hoc test. We also used ANOVAs with Dunnett’s multiple comparison test or Tukey’s post-hoc tests to assess differences in the phenotypic assays (Fig. 3). To identify differentially expressed genes in the RNA-seq data (Fig. 4), DESeq2 was used with a significance cutoff of log2 ratio of 1/-1 and FDR corrected p-value of < 0.05. For RNA-seq pathway enrichment, we used Fisher’s exact tests with Benjamini and Hochberg false discovery rate correction in the Gene-List Enrichment Tool of KOBAS. To determine if there were differences in growth across time with the different bacterial strains (Fig. 5A), a repeated-measures ANOVA was used. To assess whether *Debaryomyces* impacted the growth of *S. xylosus* BC10 ancestor and mutant strains (Fig. 5B), an ANOVA on the day 7 CFU data was used. To determine how biotic environments affected the fitness of ancestor and mutant BC10 strains (Fig. 5C), ANOVAs were used for each mutant strain. Log transformations were applied when appropriate. All statistical analyses were conducted in PRISM 9 or R.

### Data availability

All raw Illumina reads of re-sequenced evolved isolates of *S. xylosus* BC10 have been deposited in NCBI in BioProject PRJNA856679. Raw Illumina RNA-seq reads from this study have been deposited in NCBI in BioProject PRJNA856810.

## ACKNOWLEDGEMENTS

This work was funded by a grant from the United States National Science Foundation (CAREER IOS/BIO 1942063) to B.E.W. The authors are grateful to members of the Wolfe lab that provided extensive feedback on previous versions of this manuscript.

