## Supplementary material for "Species interactions promote parallel evolution of global transcriptional regulators in a widespread *Staphylococcus* species": Fig. S1

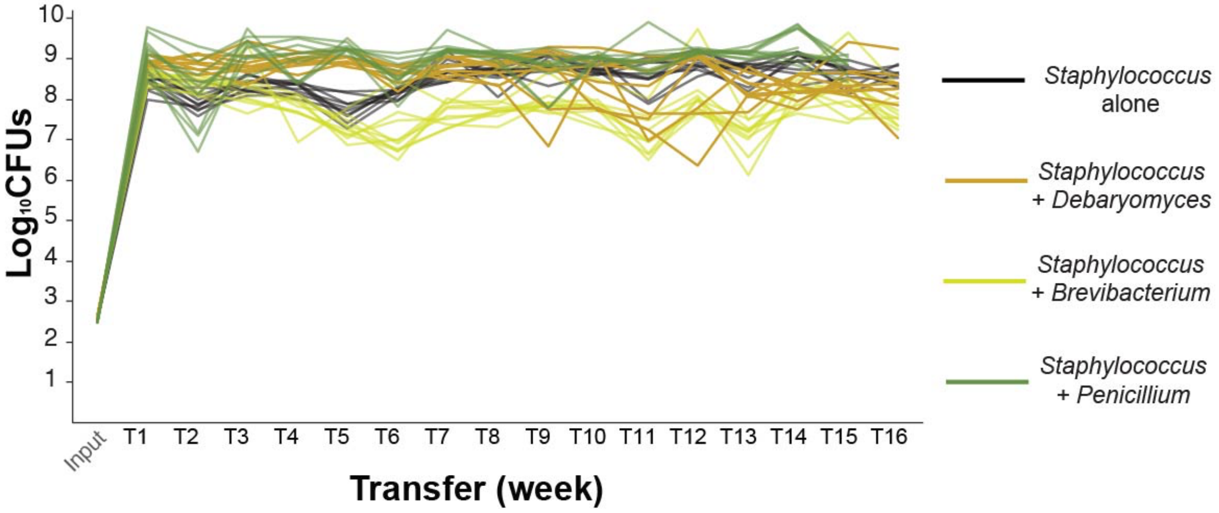


**Figure S1** - Cell densities of replicate evolution populations over time. Each line represents the total CFUs of *S. xylosus* in each experimental unit at each transfer. Because data for T16 are missing for the +*Penicillium* treatment due to a technical error, mean population size across the entire experimental evolution was calculated for T1-T15.
