## Supplementary material for "Species interactions promote parallel evolution of global transcriptional regulators in a widespread *Staphylococcus* species": Fig. S2

| | | Numbers below indicate # of isolates with non-synonymous mutation in gene | | | | | | | | | | | | | | |
|---|---|---|---|---|---|---|---|---|---|---|---|---|---|---|---|---|
| | | Alone | | | *Brevibacterium* | | | *Debaryomyces* | | | *Penicillium* | | | *Extra Debaryomyces* | | |
| Gene Annotation | Functional Category | C2 | C7 | C10 | C21 | C26 | C29 | C51 | C55 | C60 | P6 | P9 | P14 | C52 | C53 | C56 |
| FtsZ-interacting protein related to cell division | Cellular Processes | | | | 1 | | | | | | | | | | | |
| Carbon starvation protein A | Stress Response | | | | | | | | | | | | | | 1 | |
| ATP-dependent DNA helicase RecG (EC 3.6.1.-) CDS | DNA Replication | | | | | | | | | | | | | | 1 | |
| rRNA small subunit methyltransferase, glucose inhibited division protein GidB CDS | RNA Processing | | | | | | | | | | | | | | 1 | |
| Probable type II DNA modification enzyme | Defense | | 1 | | | | | | | | | | | | | |
| **Histidine kinase of the competence regulon ComD (This is actually AgrC)** | Global Regulators of Transcription | | | | | | | 4 | 1 | | | | | 1 | 1 | |
| **Response regulator of the competence regulon ComE (This is actually AgrA)** | Global Regulators of Transcription | | | | | | | 3 | | | | | | 3 | | 1 |
| **Two-component sensor kinase SA14-24 CDS (Cell wall metabolism sensor histidine kinase WalK)** | Global Regulators of Transcription | | | | | | | | | 4 | | | | 1 | 1 | 3 |
| **RsbU** | Global Regulators of Transcription | | | | | | | 1 | | | | | | | | 3 |
| **RNA polymerase sigma factor SigB CDS** | Global Regulators of Transcription | | | | | | | | 7 | 7 | | | | 3 | 2 | |
| Threonine dehydratase | Metabolism | | | 1 | | | | | | | | | | | | |
| Zinc ABC transporter, ATP-binding protein ZnuC | Metabolism | | | 1 | | | | | | | | | | | | |
| Acetate kinase | Metabolism | 1 | | | | | | | | | | | | | | |
| Hydroxymethylglutaryl-CoA synthase | Metabolism | | | | 1 | | | | | | | | | | | |
| Gluconokinase (EC 2.7.1.12) / oxidoreductase domain | Metabolism | | | | | 1 | | | | | | | | | | |
| Diaminopimelate decarboxylase (EC 4.1.1.20) CDS | Metabolism | | | | | | | | 2 | | | | | 1 | 1 | |
| Diaminopimelate epimerase alternative form predicted for S.aureus (EC 5.1.1.7) CDS | Metabolism | | | | | | | | | | | | | 1 | | |
| Glucose 1-dehydrogenase | Metabolism | | | | | | | | | | | | | 1 | | |
| Enoyl-[acyl-carrier-protein] reductase [NADH] | Metabolism | | | | | | | | | | | | | | 1 | |
| Cardiolipin synthetase (EC 2.7.8.-) | Metabolism | | | | 1 | | | | | | | | | | | |
| ABC-type Fe3+-siderophore transport system, permease 2 component CDS | Metabolism | | | | | | | | | | 1 | | | | | |
| Nicotinate-nucleotide adenylyltransferase (EC 2.7.7.18) ## bacterial NadD family CDS | Metabolism | | | | | | | | | | | | | 1 | | |
| Nicotinate phosphoribosyltransferase | Metabolism | | | | | | | | | | | | | | | 1 |
| FMN reductase (EC 1.5.1.29) | Metabolism | | | | | | | | | | | | | | 1 | |
| Peptide chain release factor 3 | Translation | | | | | 1 | | | | | | | | | | |
| CDS: tRNA-i(6)A37 methylthiotransferase | Translation | | | | | | | 1 | | | | | | | | |
| Ammonium transporter CDS | Transporters | | | | | | | | 1 | | | | | | | |
| Magnesium transporter CDS | Transporters | | | | | | | | | | | | | | 2 | |
| Membrane protein (sulfite exporter TauE/SafE family protein) | Transporters | | | | | | 1 | | | | | | | | | |
| Sodium:galactoside symporter CDS | Transporters | | | | | | | | | 1 | | | | | | |
| Oligopeptide transport ATP-binding protein OppF (TC 3.A.1.5.1) | Transporters | | | | | | | | | | | | | | 1 | |
| Oligopeptide transport ATP-binding protein OppD | Transporters | | | | | | | | | | | 2 | | | | |
| putative acyltransferase | Unknown | | | | | | | | | | | | | | 1 | |
| FIG01108899: hypothetical protein | Unknown | | | | 1 | | | | | | | | | | | |
| FIG01109350: hypothetical protein | Unknown | | | | | | 1 | | | | | | | | | |
| FIG018700 | Unknown | | | | | | | | | | 7 | | | | | |
| FIG01107972 | Unknown | | | | | | | | | | 2 | | | | | |
| Total | | 1 | 1 | 2 | 4 | 2 | 2 | 9 | 11 | 12 | 10 | 2 | 0 | 12 | 14 | 8 |
| Number of Unique | | 1 | 1 | 2 | 4 | 2 | 2 | 4 | 3 | 3 | 3 | 2 | 0 | 7 | 12 | 4 |
| % Isolates with NS SNP | | 14.29% | 14.29% | 28.57% | 42.86% | 28.57% | 28.57% | 100.00% | 100.00% | 100.00% | 100.00% | 14.29% | 0.00% | 100.00% | 100.00% | 100.00% |

**Figure S2** - Frequency of all non-synonymous mutations within a gene within a population (columns) based on sequencing 7 isolates per population. Only genes with a putative function were shown in Figure 2 of the manuscript; this figure shows all genes. It also includes the additional +*Debaryomyces* treatments that were sequenced to confirm the pattern of global regulator mutations. Numbers 1-10, 21-30, 51-60, and 61-70 are used to indicate unique populations. There were other experimental populations included in the initial experiment (31-40, 41-50), but they are excluded from this manuscript because the neighbor treatment went extinct.
