## Supplementary material for "Species interactions promote parallel evolution of global transcriptional regulators in a widespread *Staphylococcus* species": Fig. S3

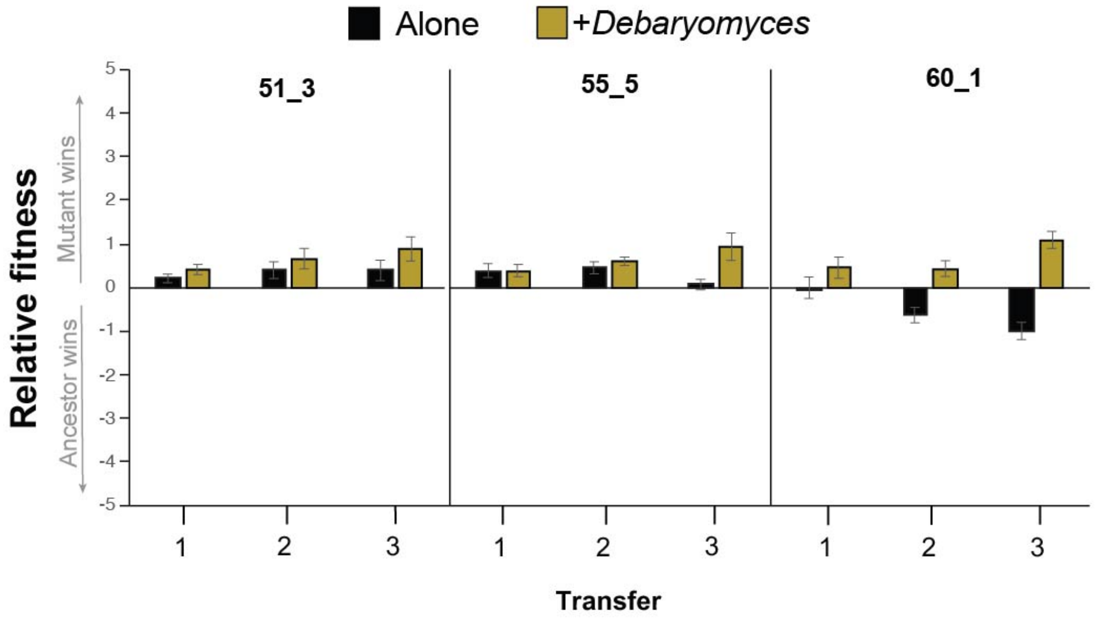


**Figure S3** - Fitness of ancestor and mutant strains of *S. xylosus* when competed in initially identical ratios either alone or in the presence of the yeast *Debaryomyces hansenii*. Relative fitness is expressed as log10((CFUs of mutant strain +1) ÷ (CFUs of ancestor strain +1)). A positive relative fitness means that the mutant strain reached a higher proportion of the total number of CFUs when competing with the ancestor. A negative fitness means the ancestor strain reached a higher proportion. Error bars represent one standard deviation of the mean. n = 8.
